# Quantitative Model of Transcriptional Noise Regulation by mRNA Condensates

**DOI:** 10.64898/2026.08.16.745099

**Authors:** Andreas Lanitis, Anatoly B. Kolomeisky

## Abstract

A fundamental biological process of transcription occurs in the cell nucleus, which is a complex medium that also contains multiple heterogeneous structures known as biomolecular condensates. Interestingly, some of these condensates contain mRNA molecules in addition to proteins, suggesting an important cellular role in transcription that is not yet well understood. In this work, we develop a minimal theoretical framework for quantitative investigation of the role of reversible mRNA condensation in transcription. Our discrete-state stochastic approach accounts for the most relevant processes, allowing us to explicitly evaluate the properties of the system and clarify the effects of condensation. Analytical calculations supported by computer simulations suggest that reversible mRNA condensation influences the transcription processes by maintaining a constant level of free mRNA in the nucleoplasm while lowering the degree of stochastic noise and increasing the robustness against external perturbations. Physicochemical arguments are presented to explain these observations. The proposed theoretical framework elucidates important microscopic aspects of transcription, providing a convenient quantitative tool for investigating complex biological phenomena.

**TOC Graphic:** 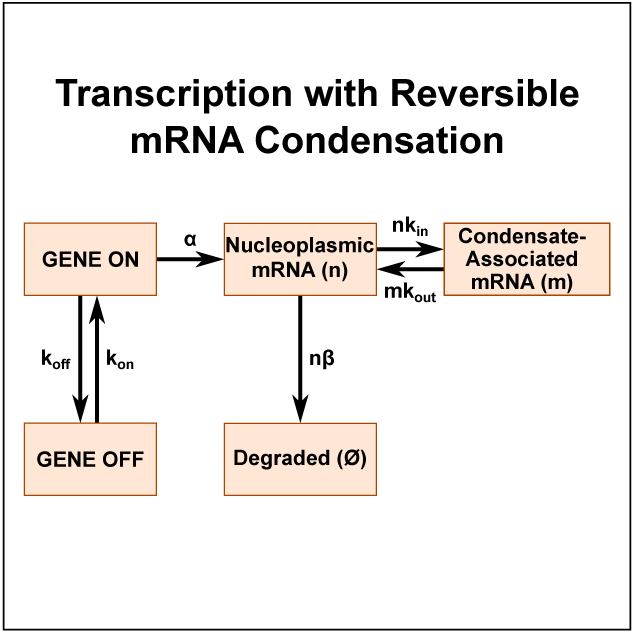

## Main

Transcription is one of the most important biological processes involved in the production of mRNA molecules by RNA polymerase enzymes which interact with the corresponding genomic regions on DNA [1–3]. It takes place in the cell nucleus, which exhibits a complex heterogeneous structure including dense liquid-like droplets known as biomolecular condensates [4–7]. These small dynamic membraneless structures allow biomolecules to freely enter, leave, associate, and dissociate as cellular conditions change [6, 8, 9]. It has been well established that some of these biomolecular condensates, such as nuclear speckles, paraspeckles, nucleoli, and Cajal bodies, contain various types of RNA molecules produced during transcription [8, 10–13]. While the important role of biomolecular condensates in regulating nuclear processes has been actively discussed [4, 5, 14, 15], many aspects of how mRNA condensates influence transcription are not fully understood [15].

To explain one of the roles of biomolecular condensates in living cells, it has been suggested theoretically via thermodynamic arguments [4, 16], also supported by some experimental observations, that liquid-liquid phase separation (LLPS), which can create such condensates, leads to the reduction of stochastic fluctuations. The main assumption is that equilibrium between dilute and dense phases buffers the concentration of biomolecules in the dilute phase against fluctuations. While the number of molecules within the condensates, the number of condensates, and their volumes fluctuate, the concentration in the dilute phase does not change due to rapid equilibration with the condensed phase. However, a significant fraction of biomolecular condensates are not equilibrium LLPS droplets [17]; instead, many of them are viscoelastic gels, dynamically arrested biomolecular assemblies or other types of non-equilibrium compartments. Nuclear condensates containing mRNA are likely to be non-equilibrium systems due to multiple active energy-dissipation processes, such as transcription, splicing and ATP-dependent remodeling, occurring in a small nuclear volume [18].

The role of RNA molecules in nuclear condensates has recently been discussed in the context of an RNA-mediated feedback control mechanism of transcription [19, 20]. RNA molecules in small amounts have been argued to stimulate the formation of transcriptional condensates that help transcription factors bind more strongly to their specific sites in DNA, while the production of too many RNA molecules destabilizes the condensates, stopping transcription. This elegant theoretical idea essentially explains the start and end of a transcriptional burst, a period of continuous production of mRNA, via a non-equilibrium feedback mechanism that relies on these molecules. However, it remains unclear what the role of mRNA condensates is during a transcriptional burst.

Currently, there is a very limited understanding of how mRNA condensates might affect the dynamics of transcription [21, 22]. The goal of our work is to fill this knowledge gap. We present a minimal theoretical model that couples the processes of mRNA production, degradation, and reversible condensation by applying a non-equilibrium discrete-state stochastic approach that can be explicitly analyzed. It allows us to obtain a fully quantitative description of transcriptional dynamics in the presence of reversible mRNA condensation via analytical calculations and extensive Monte Carlo computer simulations. Our analysis shows that mRNA condensates strongly influence the dynamics of transcription. They help maintain a stable mean number of free mRNA molecules in the nucleoplasm, decrease the degree of stochastic fluctuations, and improve the robustness of the transcriptional system against fluctuations in mRNA concentrations, leading to substantially reduced, though still super-Poissonian, noise.

It is important to note that our theoretical method differs from the recent model that investigated noise reduction by mRNA assembly [21]. Their model attempts to describe mRNA condensates formed through liquid-liquid phase separation, where the mRNA molecules are directly involved in their formation. However, as already discussed, not every condensate is formed through the same mechanism and mRNA molecules do not always play a key role in the formation of the condensates. Our work assumes preformed condensates, independently of their mechanism of formation, and therefore applies to any mRNA system in which mRNA molecules are not fully responsible for condensate formation. We provide a closed-form, exactly solvable master-equations framework, which leads to a clearer picture of the underlying processes.

Let us consider transcription inside the cell nucleus in the presence of mRNA condensates, as illustrated in Fig. 1. RNA polymerases move along the genomic region of DNA and synthesize mRNA molecules with rate *α* if the gene is in the active ON state. The produced mRNA species can each be degraded or removed from the cell nucleus with a rate constant *β*. This means that the total degradation rate is *nβ* if at a given time there are *n* free mRNA molecules in the system. The gene can transition from the active transcription state ON to the inactive state OFF with rate *k_off_*, while the reverse activation transition occurs with rate *k_on_*: see Fig. 1. In the OFF state, mRNA molecules are no longer produced, while degradation still takes place with the same rate constant *β*. In our model, *n* represents the number of free mRNA molecules, while we define the number of condensate-associated mRNA molecules as *m*. Free mRNA molecules can individually transition into the biomolecular condensates with rate constant *k_in_*, while condensate-associated mRNA molecules can dissociate with rate constant *k_out_*. Thus, the total rates for reversible mRNA condensation are *nk_in_* and *mk_out_*, respectively. The model is non-equilibrium, and no assumptions of equilibrium are made a priori. To simplify calculations and better portray the effects of reversible mRNA condensation on transcription, we further assume that condensate-associated mRNA molecules are not degraded. However, we have extended the model to incorporate this process and we present analytical results and simulations in the Supporting Information. Although accounting for this effect alters the mean numbers of mRNA molecules in the nucleoplasm and the condensates, it does not change the main predictions of our analysis using the minimal model.

**Figure 1:**
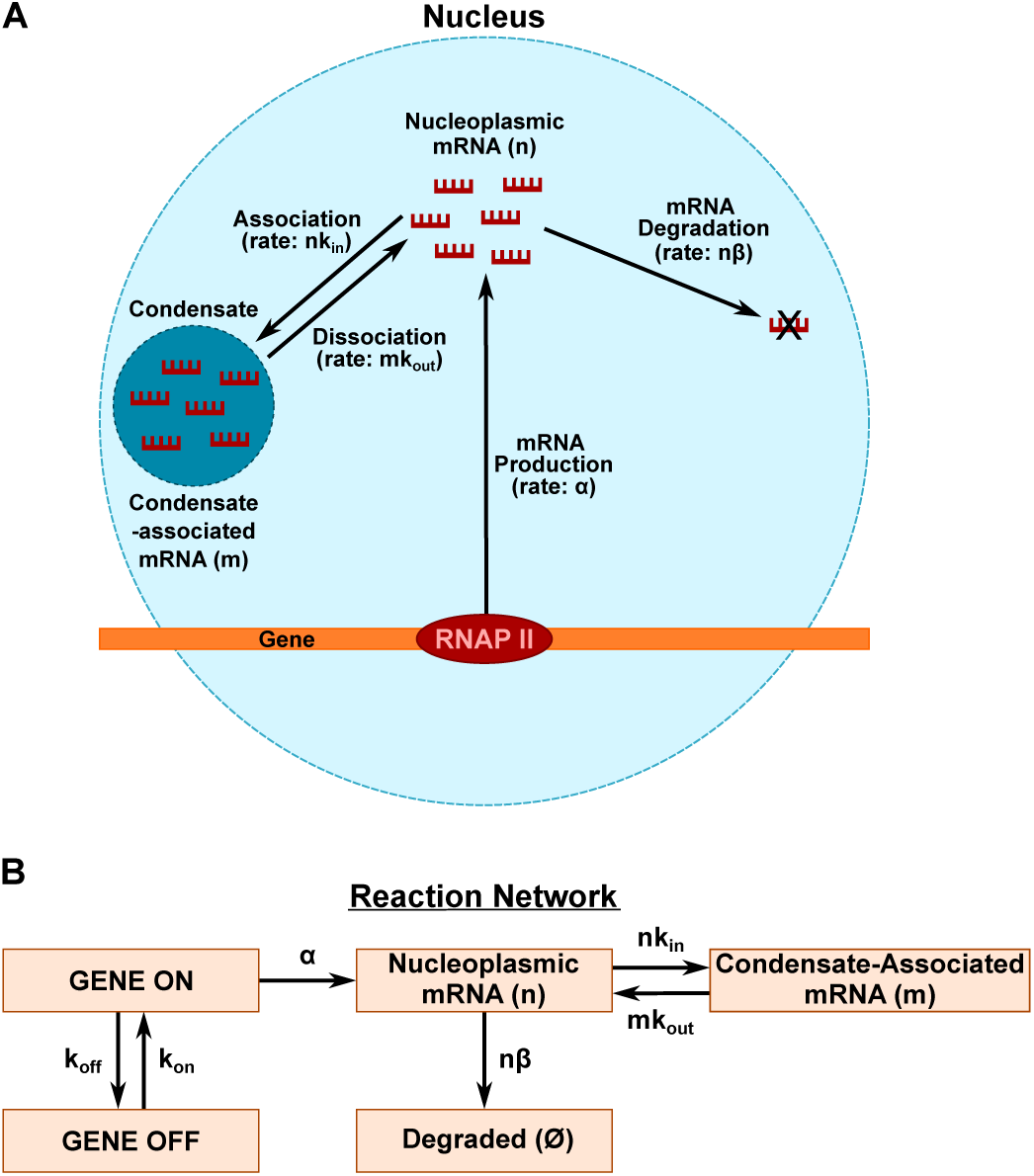
Schematic view of transcription with reversible mRNA condensation. A) Details of processes in the cell nucleus. RNAP enzymes move along the DNA chain and catalyze the production of mRNA molecules if the gene is activated (ON state). The synthesized mRNA molecules are degraded or removed from the cell nucleus when the gene is activated or deactivated. The produced mRNA molecules are distributed between two phases: free in the nucleoplasm or inside the biomolecular condensates. B) Details of stochastic processes in the system.

The main idea of our theoretical approach is to extend the original discrete-state stochastic model that was able to explicitly describe transcriptional bursting [23, 24] by coupling it with the reversible exchange of mRNA molecules between the nucleoplasm and biomolecular condensates. The corresponding kinetic scheme for stochastic transitions in the system is presented in Fig. 2. We define 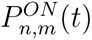 and 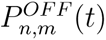 as the probabilities of finding the system in the state with *n* free mRNA molecules in the nucleoplasm and *m* mRNA molecules in the biomolecular condensates at time *t* if the gene is in the *ON* or *OFF* states, respectively. These probabilities are normalized by the condition 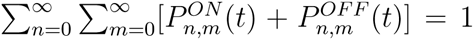. Their temporal evolution for the interior states (*n, m >* 0) follows the set of forward master equations,

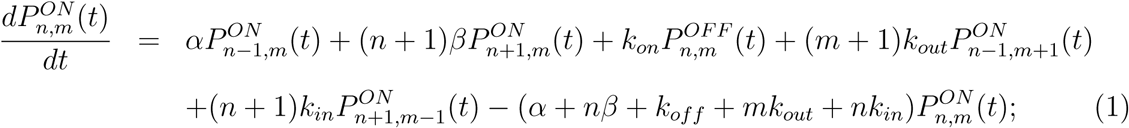

**Figure 2:**
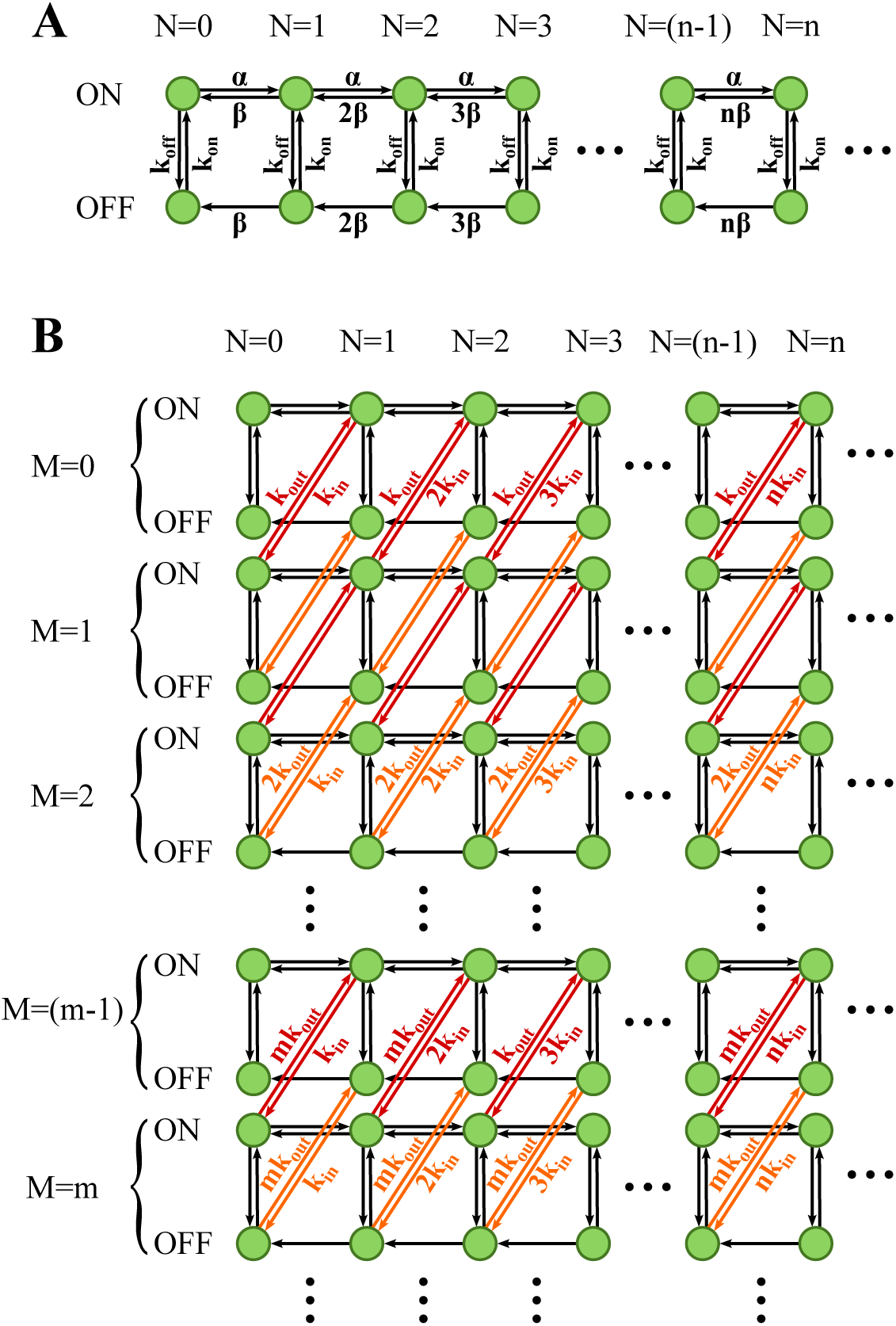
Schematic representations of the stochastic network describing transcription in the presence of condensates. A) The two-state model of transcriptional bursting without condensation. B) The two-state model with reversible mRNA exchange between the nucleoplasm and the condensates. Black arrows correspond to the same transitions as in the two-state model, while red and orange arrows correspond to mRNA exchange transitions when the gene is in the ON or OFF state, respectively. More details of the model are given in the text.

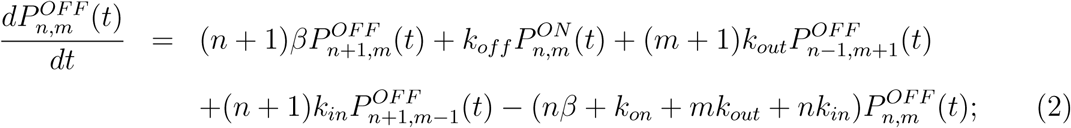

The complete set of master equations, including boundary conditions, is shown in the Supporting Information. There, they are analytically solved using the method of generating functions [23, 25], allowing us to obtain quantitative results for the stationary state (*t* → ∞). More specifically, we are interested in the mean number of mRNA molecules in the nucleoplasm, ⟨*n*⟩, and the mean number of mRNA molecules in the condensates, ⟨*m*⟩, defined as

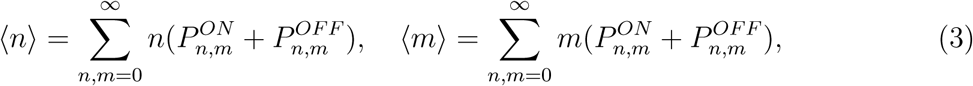

where 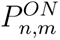 and 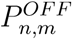 are stationary probabilities.

The explicit calculations, presented in the Supporting Information, produce the following formulas,

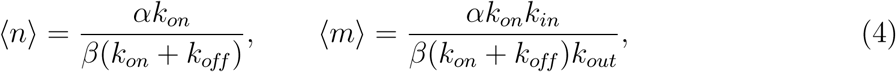

which immediately lead to

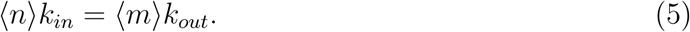

This expression can be explained using the following arguments. The overall condensation dynamics in the system can be viewed as mRNA transitions between two macrostates, one corresponding to free mRNA molecules in the nucleoplasm, and another one corresponding to all other mRNA molecules in the biomolecular condensates. Since there are only two macrostates, in the stationary limit an effective balance will be reached between them, which is reflected in Eq. 5. Another way to see it is to notice that condensation only redistributes mRNA between the two pools and is fully reversible, while the amount of free mRNA is directly controlled by synthesis and degradation. Then, under steady-state conditions, the net condensation flux must vanish, which means that the condensation will not influence the balance between mRNA production and degradation.

The results of our explicit calculations of the mean numbers of mRNA molecules in different phases of the nucleoplasm, supplemented by Monte Carlo computer simulations, are presented in Fig. 3. As expected, increasing the production rate *α* (when all other rates are fixed) leads to larger mean numbers of mRNA molecules, both in the nucleoplasm and in the condensates (Figs. 3A and 3B). Similarly, reducing the frequency of gene deactivation (smaller *k_off_*) produces larger amounts of mRNA molecules in both phases (Figs. 3C and 3D). This is also the expected result because increasing *α* leads to stronger mRNA production during transcriptional bursts, with all other parameters fixed, and decreasing *k_off_* leads to longer burst durations. Since these changes increase the mRNA production flux, balance must be achieved at higher mean mRNA values. It is also important to note that in our calculations we used experimentally relevant transition rates, and our estimates of free mRNA molecules, ranging from 5 to 50 molecules per gene, are of the same order of magnitude as available experimental estimates [26].

**Figure 3:**
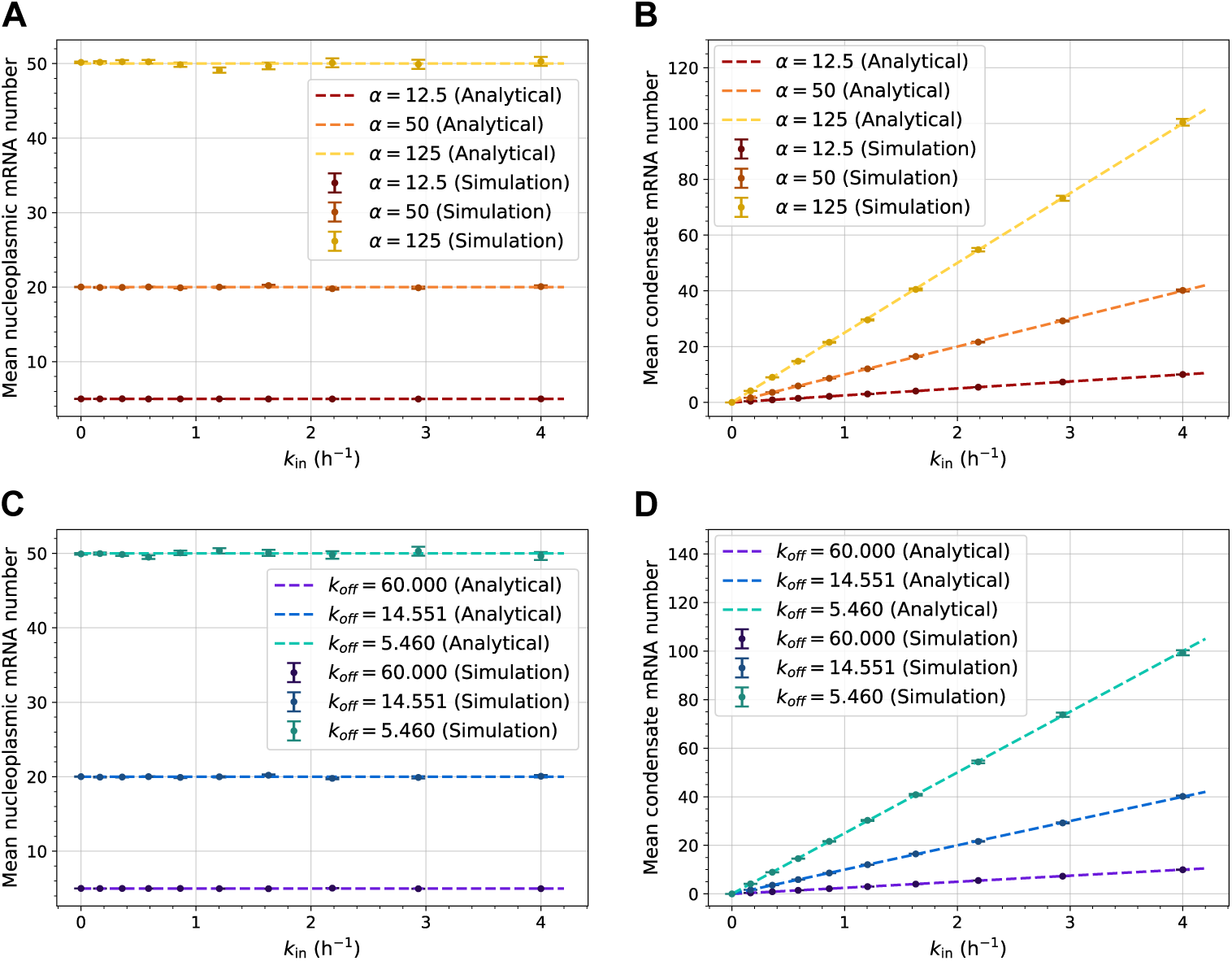
Mean numbers of mRNA molecules in different phases of the nucleoplasm as a function of the tendency to transition into the condensates. A) Mean number of free mRNA molecules ⟨*n*⟩ for different mRNA production rates. B) Mean number of condensed mRNA molecules ⟨*m*⟩ for different mRNA production rates. C) Mean number of free mRNA molecules ⟨*n*⟩ for different gene deactivation rates. D) Mean number of condensed mRNA molecules ⟨*m*⟩ for different gene deactivation rates. Lines are analytical predictions, and symbols are from Monte Carlo computer simulations. In all plots, *k_in_* is treated as an effective measure of mRNA condensation and the error bars represent the standard errors of the simulated values, calculated by block averaging (see computer simulation details in the Supporting Information).

However, the most surprising observation from our analysis is that while the number of mRNA in condensates, ⟨*m*⟩, increases with a stronger tendency to condense (Figs. 3B and 3D), the amount of free mRNA molecules, ⟨*n*⟩, is not affected at all by condensation (Figs. 3A and 3C). This can be explained by noting, as already discussed, that the stationary-state balance for ⟨*n*⟩ is not affected by condensation. At the same time, the amount of mRNA in condensates increases proportionally to the condensation rate constant *k_in_* (for fixed *k_out_*). The ratio *k_in_/k_out_* can be viewed as an effective “equilibrium” constant of condensation (although the system is still out of equilibrium), and a stronger tendency for mRNA molecules to condense is expected for larger *k_in_*. This is an important result, indicating that mRNA condensation does not affect the amount of free mRNA molecules that should eventually go to the cytoplasm to participate in the production of corresponding proteins. Thus, the presence of mRNA condensates does not decrease the overall efficiency of transcription and downstream translation processes.

The advantage of our theoretical method is that it can explicitly evaluate the degree of stochastic fluctuations for mRNA molecules in the nucleoplasm, quantifying the noise in the system. It is convenient to express it using a dimensionless parameter *F*, known as the Fano factor [23, 27], which already accounts for the changes in the mean values when quantifying stochastic fluctuations. It is defined as

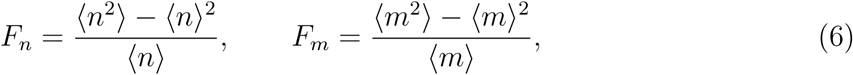

for the free mRNA and for the mRNA in the condensates, respectively. In the limit where the overall transcription can be described as a Poisson single-state process, we have *F_n_* = 1 and *F_m_* = 1, while for a more realistic system with multiple transitions and reversible condensation it is expected that *F_n/m_ >* 1. The larger the Fano factor, the larger the stochastic noise in the system. Since our theoretical method can exactly calculate the first two moments of the mRNA copy-number distributions, as explained in detail in the Supporting Information, we can obtain the explicit expressions,

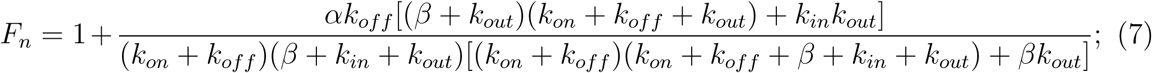

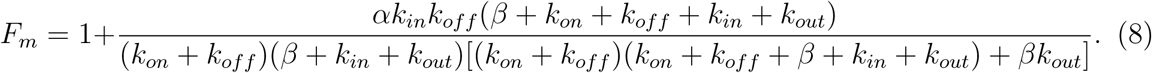

Note that in the limit of no condensation (*k_in_* → 0 or *k_out_* → ∞), the expression for *F_n_*reduces to the known result of the Fano factor for the two-state transcriptional bursting model without condensation (free mRNA molecules) [23].

The results of our analytical predictions and computer simulations for the Fano factor at different ranges of transition rates are presented in Fig. 4. Increasing the mRNA production rate *α* or decreasing the gene deactivation rate *k_off_* leads to a larger degree of stochastic noise in the system (Fig. 4). This can be understood through a simplified physical picture of a typical fluctuation. In the bursty regime, we can qualitatively represent fluctuations as cycles between minimum and maximum values around the mean. Starting near the minimum, the system transitions into the genomic ON state, where a transcriptional burst with average size *α/k_off_* drives the mRNA number towards the maximum. The gene then transitions to the OFF state, during which degradation drives the system back toward the minimum. For *β/k_on_* ≪ 1, the characteristic decrease during an OFF period can be approximated as *nβ/k_on_*. Since the upward and downward excursions must balance over a typical cycle, *n* scales approximately as *α/k_off_* as well. Thus, the characteristic fluctuation amplitude Δ*n* scales linearly with *n*, rather than with 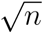 as in Poissonian statistics. We therefore expect fluctuations to become increasingly super-Poissonian, as the production rate *α* increases or the frequency of deactivation *k_off_* decreases, consistent with Fig. 4. We state that this picture is intended only as a qualitative illustration of the dependence of stochastic noise on the kinetic parameters. One could also note that at low values of ⟨*n*⟩ and ⟨*m*⟩, the lower boundary at zero additionally limits fluctuations, since *n*, *m* cannot become negative.

**Figure 4:**
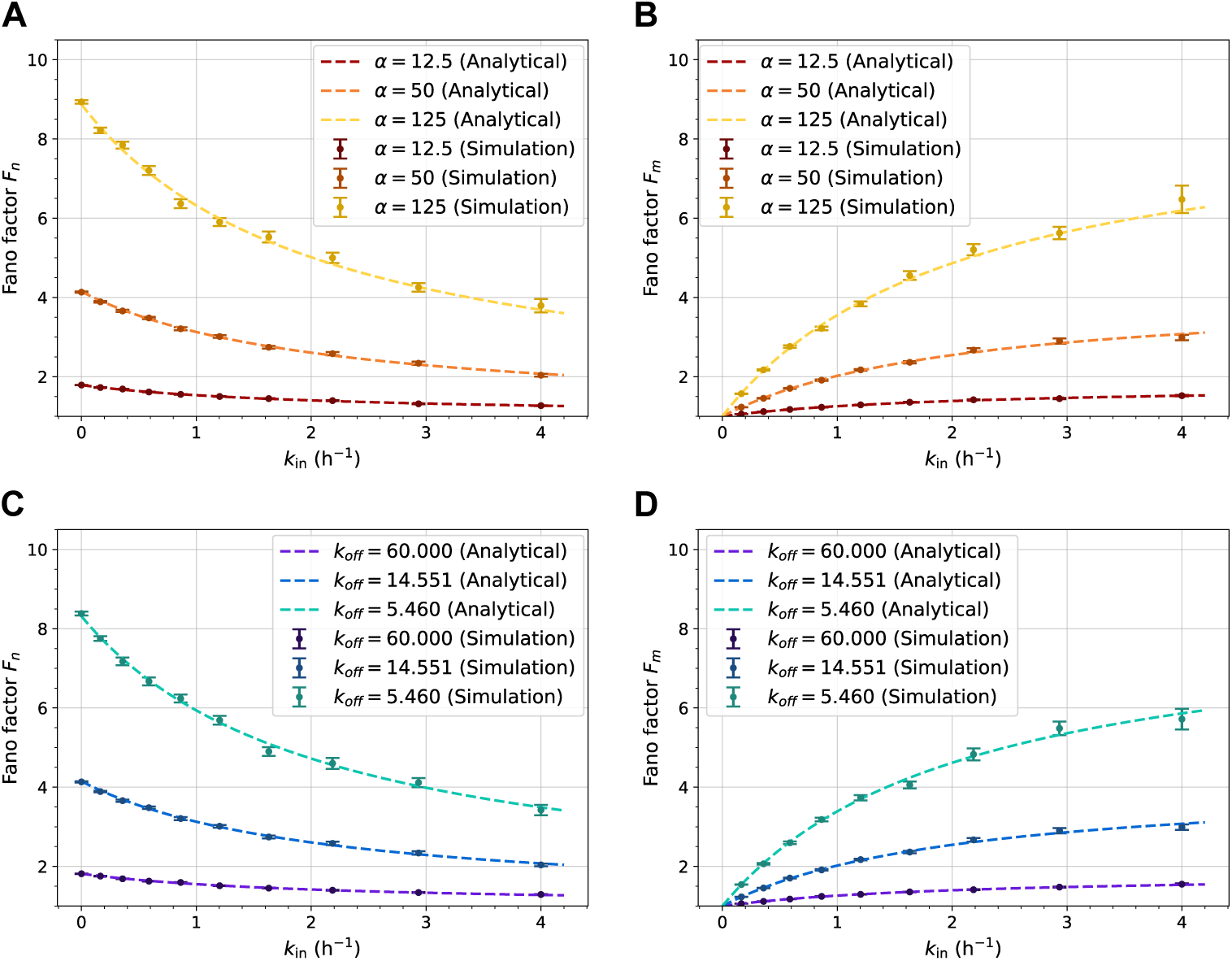
Degree of stochastic fluctuations, quantified by Fano factors, for mRNA molecules in different phases of the nucleoplasm as a function of the tendency to transition into the condensates. A) Fano factor of free mRNA molecules, *F_n_*, for different mRNA production rates. B) Fano factor of condensed mRNA molecules, *F_m_*, for different mRNA production rates. C) Fano factor of free mRNA molecules, *F_n_*, for different gene deactivation rates. D) Fano factor of condensed mRNA molecules, *F_m_*, for different gene deactivation rates. Lines are analytical predictions, and symbols are from Monte Carlo computer simulations. In all plots, *k_in_* is treated as an effective measure of mRNA condensation and the error bars represent the standard errors of the simulated values, calculated by block averaging (see computer simulation details in the Supporting Information).

Observing Fig. 4 with regard to the tendency of mRNA molecules to condense, we see that condensation lowers the stochastic noise for free mRNA molecules, while it simultaneously increases the degree of stochastic fluctuations for mRNA molecules in the condensates. This can be understood by identifying mRNA condensates as mRNA reservoirs, buffering against stochastic fluctuations in the free mRNA, while *k_in_* acts as the coupling strength between them. As the free mRNA population displays fluctuations around the stationary state, there exists a restoring exchange current between the free mRNA pool and the reservoir, which becomes increasingly strong as *k_in_* increases. Thus, as the tendency to condense increases, free mRNA noise is increasingly suppressed, while the larger transfers of mRNA between the condensates and the nucleoplasm cause larger fluctuations in the condensates. The decrease in nucleoplasmic mRNA noise is observed for all kinetic parameters that govern the transitions in the system, and we conclude that noise suppression is an important function of biomolecular condensates during transcription.

The cell nucleus is a complex dynamic system that frequently experiences large perturbations in biomolecular composition. It is important to understand how these fluctuations will affect transcription to further elucidate the role of mRNA condensates. One can explicitly investigate this question using our theoretical model. For this purpose, we used stochastic simulations to investigate the mean first-passage time to the stationary nucleoplasmic mean mRNA number ⟨*n*⟩ after an instantaneous change in the number of free mRNA molecules. To do so, we initiated computer simulations of the stochastic network at initial states (⟨*n*⟩ + *x,* ⟨*m*⟩), corresponding to the addition (*x >* 0) or removal (*x <* 0) of |*x*| mRNA molecules from the system at *t* = 0. The initial promoter state was selected based on the occupation probabilities *P^ON^* and *P^OF^^F^*. We thus measured the mean first-passage times for a range of values of *k_in_* and *x* = ±2, ±4, as presented in Fig. 5A. A monotonic decrease in mean first-passage times is observed as the tendency to condense increases (larger *k_in_*), which suggests that another important function of mRNA condensation is to make transcription a more robust process, helping to quickly neutralize the effects of sudden perturbations. In Fig. 5A, the mean first-passage times for *x* = −2, −4 are initially larger than those for *x* = +2, +4, since they require a transcriptional burst to reach the stationary state, whereas positive perturbations can relax directly through degradation. However, as the value of *k_in_*increases, the drive towards the stationary state due to the condensates begins to dominate, eventually causing the mean first-passage times for *x* = ±2 and *x* = ±4 to converge. In this regime, only the distance from the stationary state affects the mean first-passage time and not the bursty nature of transcription. This suggests that biomolecular condensates serve to maintain the robustness of transcription, without relying directly on transcriptional bursts for relaxation.

**Figure 5:**
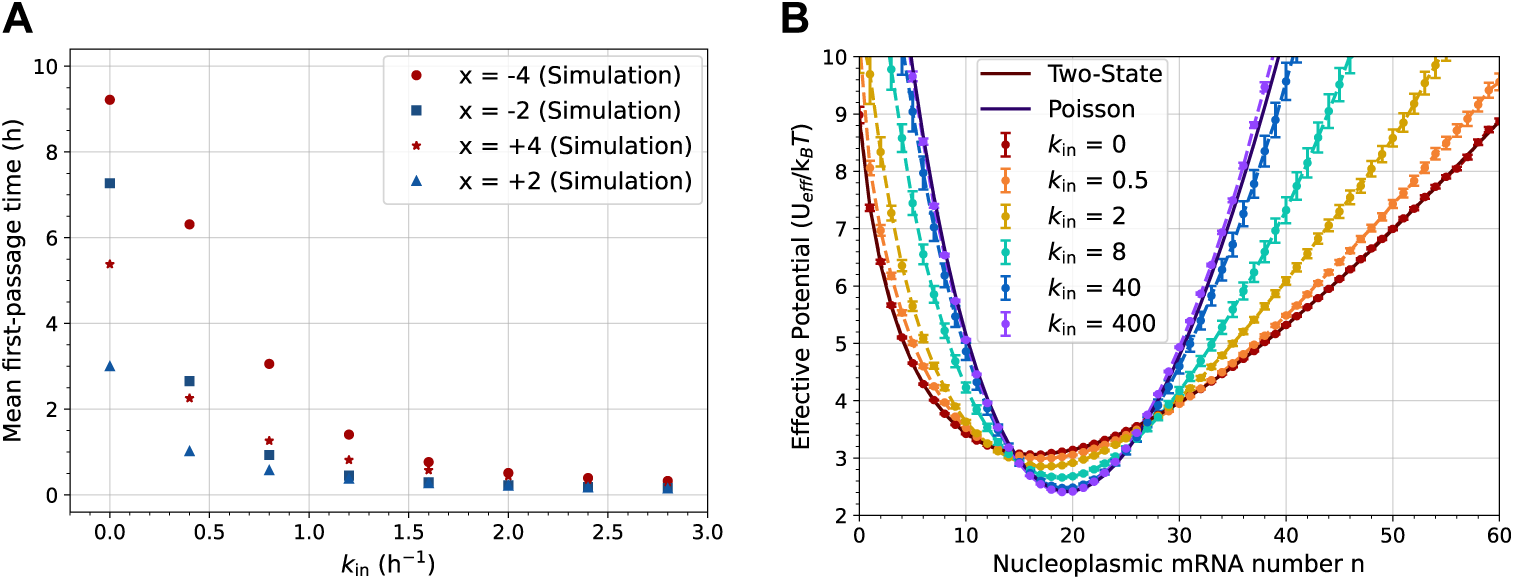
Relaxation dynamics to stationary behavior after a sudden perturbation in the number of free mRNA molecules. A) Mean first-passage times to return to the mean nucleoplasmic mRNA number ⟨*n*⟩ for simulations initiated at *t* = 0 in the state [⟨*n*⟩ + *x*, ⟨*m*⟩] versus *k_in_*. B) Effective potentials illustrating the relaxation dynamics. The potentials are estimated using Eq. 9 via computer simulations or analytical calculations. In both graphs, *k_in_* is treated as an effective measure of mRNA condensation and the error bars represent the standard errors of the simulated values (simulation and analysis details in Supporting Information and the main text).

To better illustrate the faster relaxation dynamics for the system with increased mRNA condensation, we introduce the idea of an effective potential, which can be defined as

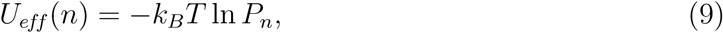

where 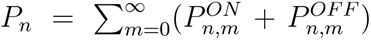 is the stationary probability of observing *n* mRNA molecules in the nucleoplasm. The physical meaning of this effective potential can be understood through the following simplified qualitative picture. In the stationary limit, the most probable value of *n* is close to ⟨*n*⟩ (but generally not the same since the potential might not be symmetric), and the system fluctuates around this point. Qualitatively, the dynamics of the system can be viewed as the motion of a fictitious particle in the effective potential *U_eff_* (*n*). Within this simplified picture, a steeper potential corresponds to a stronger restoring tendency toward the stationary point and is therefore associated with faster relaxation dynamics. This effective-potential approach provides an intuitive illustration of the role of mRNA condensates in relaxation dynamics after perturbations.

We have estimated the effective potential computationally for our model of transcription with reversible mRNA condensation using a range of *k_in_* values and analytically for the limiting cases of the two-state model without condensation [25, 28] (as shown in Fig. 2A) and for the Poisson single-state limit [23] (no gene deactivation), and the details are presented in the Supporting Information. The results of our calculations and computer simulations are presented in Fig. 5B. In the limit *k_in_* → 0, when our model displays no mRNA condensation, the computer simulations fully agree with the analytical result for the two-state transcriptional bursting model without condensation. Increasing the tendency to condense (larger *k_in_*) makes the effective potential steeper, approaching the analytical Poisson limit for *k_in_* ≫ 1. This steepening is consistent with the faster return to stationary values after sudden perturbations observed in Fig. 5A. Note that the effective potential is not symmetric about the most probable value of *n*, reflecting the fact that the sudden perturbations that remove or add the same number of free mRNA molecules do not display the same mean first-passage time: compare the curves for *x >* 0 and *x <* 0 in Fig. 5A. However, it becomes increasingly symmetric as it approaches the Poisson single-state limit. Overall, the effective potential conveniently illustrates the response of the system to perturbations, suggesting that with increasing condensation transcription becomes more robust.

Throughout the study, we performed extensive Monte Carlo computer simulations of the stochastic network for transcription with mRNA condensation using the Gillespie algorithm. We selected biologically motivated rates for production, degradation, gene activation and deactivation, based on measurements conducted in mouse fibroblasts [29], without attempting to model a specific gene. The condensation and dissociation rate constants *k_in_* and *k_out_*were instead treated as model parameters. Tables containing the exact kinetic rates used in each simulation are presented in the Supporting Information.

Although our theoretical method was able to quantitatively describe transcription in the presence of reversible mRNA condensation and obtained analytical results for some quantities that are consistent with experimental estimates, it is important to discuss its limitations. In our approach, it was implicitly assumed that the condensates constitute one continuous phase, which allowed us to assume fixed rate constants for entry into and exit from the condensate. A more realistic picture would be that there are multiple liquid-like droplets of biomolecular condensates with variable amounts of mRNA molecules, which would lead to distributions of the association and dissociation rates. Thus, our picture is rather a simplified homogeneous mean-field description of the complex heterogeneous process of condensation. In addition, many more biochemical states and reactions are involved in mRNA production and degradation, which are not accounted for in our approach [23, 30, 31]. Furthermore, it is assumed that the initiation and termination of a transcriptional burst are independent of mRNA molecules, which might not be fully in line with recent experimental studies [19, 20]. However, despite these limitations, the proposed theoretical framework clarifies important mechanistic aspects of transcription and condensation, providing a convenient method to quantitatively evaluate their complex biological processes.

In conclusion, we developed a minimal theoretical approach to investigate the role of reversible mRNA condensation in the cell nucleus during transcription. Specifically, our model views transcription as a two-state process, with mRNA being synthesized in the genomic ON state and being degraded in both activated and deactivated states. Furthermore, mRNA molecules can enter and exit the condensed phase depending on the relative numbers of mRNA molecules in both phases. The corresponding discrete-state stochastic model was solved analytically using the method of generating functions, allowing us to estimate the properties of transcription in the stationary limit. Our analytical calculations were also supported by Monte Carlo computer simulations. We found that reversible condensation plays a crucial role in transcription. The ability of mRNA molecules to condense maintains the mean number of free mRNA molecules in the cell nucleus, reduces stochastic noise and makes the system more robust against sudden perturbations. Thus, we argue that mRNA condensates act as a powerful mechanism for quickly and efficiently regulating transcription. The proposed mechanism might also be applicable, with appropriate extensions, to mRNA in biomolecular condensates outside of the cell nucleus and to proteins. Importantly, our theoretical method provides quantitative predictions that might be directly tested in experiments, opening new directions and approaches for investigating complex biological phenomena.

## Supporting Information

Details of analytical calculations using the generating functions method, and details of the Monte Carlo simulations scheme.

## Acknowledgment

We acknowledge the support by the Welch Foundation (C-1559), and by the Center for Theoretical Biological Physics sponsored by the NSF (PHY-2019745).

