## Supporting Information for "Quantitative Model of Transcriptional Noise Regulation by mRNA Condensates"

### Supporting Information: Quantitative Model of Transcriptional Noise Regulation by mRNA Condensates

### Analytical Derivations

#### Extended model without condensate-associated mRNA degradation

The extended model network with no condensate-associated degradation is described by the master equations:

$\underline{N = 0,\ M = 0}$:

$$\frac{dP_{00}^{on}}{dt} = k_{on}P_{00}^{off} + \beta P_{10}^{on} - (k_{off} + \alpha)P_{00}^{on} \tag{S1}$$

$$\frac{dP_{00}^{off}}{dt} = k_{off}P_{00}^{on} + \beta P_{10}^{off} - k_{on}P_{00}^{off} \tag{S2}$$

$\underline{N \neq 0,\ M = 0}$:

$$\begin{aligned}\frac{dP_{n0}^{on}}{dt} &= k_{on}P_{n0}^{off} + k_{out}P_{(n-1)1}^{on} + (n+1)\beta P_{(n+1)0}^{on} + \alpha P_{(n-1)0}^{on} \\ &\quad - (k_{off} + n\beta + nk_{in} + \alpha)P_{n0}^{on}\end{aligned} \tag{S3}$$

$$\frac{dP_{n0}^{off}}{dt} = k_{off}P_{n0}^{on} + k_{out}P_{(n-1)1}^{off} + (n+1)\beta P_{(n+1)0}^{off} - (k_{on} + n\beta + nk_{in})P_{n0}^{off} \tag{S4}$$

$\underline{N = 0,\ M \neq 0}$:

$$\frac{dP_{0m}^{on}}{dt} = k_{on}P_{0m}^{off} + k_{in}P_{1(m-1)}^{on} + \beta P_{1m}^{on} - (k_{off} + mk_{out} + \alpha)P_{0m}^{on} \tag{S5}$$

$$\frac{dP_{0m}^{off}}{dt} = k_{off}P_{0m}^{on} + k_{in}P_{1(m-1)}^{off} + \beta P_{1m}^{off} - (k_{on} + mk_{out})P_{0m}^{off} \tag{S6}$$

$\underline{N \neq 0,\ M \neq 0}$:

$$\frac{dP^{on}_{nm}}{dt} = k_{on}P^{off}_{nm} + (n+1)k_{in}P^{on}_{(n+1)(m-1)} + (m+1)k_{out}P^{on}_{(n-1)(m+1)} + (n+1)\beta P^{on}_{(n+1)m} + \alpha P^{on}_{(n-1)m} - (k_{off} + nk_{in} + mk_{out} + n\beta + \alpha)P^{on}_{nm} \tag{S7}$$

$$\frac{dP^{off}_{nm}}{dt} = k_{off}P^{on}_{nm} + (n+1)k_{in}P^{off}_{(n+1)(m-1)} + (m+1)k_{out}P^{off}_{(n-1)(m+1)} + (n+1)\beta P^{off}_{(n+1)m} - (k_{on} + nk_{in} + mk_{out} + n\beta)P^{off}_{nm} \tag{S8}$$

To derive analytical results, we use the generating functions:

$$G^{on}(x,y,t) = \sum_{n,m=0}^{\infty} x^n y^m P^{on}_{nm}(t) \qquad G^{off}(x,y,t) = \sum_{n,m=0}^{\infty} x^n y^m P^{off}_{nm}(t) \tag{S9}$$

Transforming the master equations gives:

$$\frac{\partial G^{on}}{\partial t} = k_{on}G^{off} - k_{off}G^{on} + \alpha(x-1)G^{on} + \beta(1-x)\frac{\partial G^{on}}{\partial x} + k_{out}(x-y)\frac{\partial G^{on}}{\partial y} + k_{in}(y-x)\frac{\partial G^{on}}{\partial x} \tag{S10}$$

$$\frac{\partial G^{off}}{\partial t} = k_{off}G^{on} - k_{on}G^{off} + \beta(1-x)\frac{\partial G^{off}}{\partial x} + k_{out}(x-y)\frac{\partial G^{off}}{\partial y} + k_{in}(y-x)\frac{\partial G^{off}}{\partial x} \tag{S11}$$

We can derive analytical results from the following relations:

$$G^{on/off}(1,1,t) = \sum_{n,m=0}^{\infty} P_{nm}^{on/off}(t) = P^{on/off}(t) \tag{S12}$$

$$\left.\frac{\partial G^{on/off}}{\partial x}\right|_{\substack{x=1\\y=1}} = \left.\sum_{n,m=0}^{\infty} nx^{n-1}y^m P_{nm}^{on/off}(t)\right|_{\substack{x=1\\y=1}} = \sum_{n,m=0}^{\infty} nP_{nm}^{on/off}(t) = \langle n(t)\rangle^{on/off} \tag{S13}$$

$$\left.\frac{\partial G^{on/off}}{\partial y}\right|_{\substack{x=1\\y=1}} = \left.\sum_{n,m=0}^{\infty} mx^{n}y^{m-1} P_{nm}^{on/off}(t)\right|_{\substack{x=1\\y=1}} = \sum_{n,m=0}^{\infty} mP_{nm}^{on/off}(t) = \langle m(t)\rangle^{on/off} \tag{S14}$$

$$\begin{aligned}\left.\frac{\partial^2 G^{on/off}}{\partial x \partial y}\right|_{\substack{x=1\\y=1}} &= \left.\sum_{n,m=0}^{\infty} nmx^{n-1}y^{m-1} P_{nm}^{on/off}(t)\right|_{\substack{x=1\\y=1}} = \sum_{n,m=0}^{\infty} nmP_{nm}^{on/off}(t) \\ &= \langle nm(t)\rangle^{on/off}\end{aligned} \tag{S15}$$

$$\begin{aligned}\left.\frac{\partial^2 G^{on/off}}{\partial x^2}\right|_{\substack{x=1\\y=1}} &= \left.\sum_{n,m=0}^{\infty} n(n-1)x^{n-2}y^{m} P_{nm}^{on/off}(t)\right|_{\substack{x=1\\y=1}} = \sum_{n,m=0}^{\infty} (n^2-n)P_{nm}^{on/off}(t) \\ &= \langle n^2(t)\rangle^{on/off} - \langle n(t)\rangle^{on/off}\end{aligned} \tag{S16}$$

$$\begin{aligned}\left.\frac{\partial^2 G^{on/off}}{\partial y^2}\right|_{\substack{x=1\\y=1}} &= \left.\sum_{n,m=0}^{\infty} m(m-1)x^{n}y^{m-2} P_{nm}^{on/off}(t)\right|_{\substack{x=1\\y=1}} = \sum_{n,m=0}^{\infty} (m^2-m)P_{nm}^{on/off}(t) \\ &= \langle m^2(t)\rangle^{on/off} - \langle m(t)\rangle^{on/off}\end{aligned} \tag{S17}$$

Setting $x = 1$, $y = 1$ in equations S10 and S11:

$$\frac{dP^{on}(t)}{dt} = k_{on}P^{off}(t) - k_{off}P^{on}(t) \qquad \frac{dP^{off}(t)}{dt} = k_{off}P^{on}(t) - k_{on}P^{off}(t) \tag{S18}$$

At the steady state ($\frac{dP^{on/off}}{dt} = 0$), with the condition $P^{on} + P^{off} = 1$, we derive:

$$P^{on} = \frac{k_{on}}{k_{on}+k_{off}} \qquad P^{off} = \frac{k_{off}}{k_{on}+k_{off}} \tag{S19}$$

Since $\langle n(t)\rangle = \langle n(t)\rangle^{on} + \langle n(t)\rangle^{off}$ and $\langle m(t)\rangle = \langle m(t)\rangle^{on} + \langle m(t)\rangle^{off}$, we add together the partial derivatives of equations S10 and S11 with respect to x for $x = 1$, $y = 1$, and with respect to y for $x = 1$, $y = 1$:

$$\frac{d\langle n\rangle}{dt} = \alpha P^{on} - \beta\langle n\rangle - k_{in}\langle n\rangle + k_{out}\langle m\rangle \tag{S20}$$

$$\frac{d\langle m\rangle}{dt} = k_{in}\langle n\rangle - k_{out}\langle m\rangle \tag{S21}$$

Solving the system for the steady state, using $P^{on}$ gives:

$$\langle n\rangle = \frac{\alpha k_{on}}{\beta(k_{on} + k_{off})} \qquad \langle m\rangle = \frac{k_{in}}{k_{out}}\langle n\rangle = \frac{\alpha k_{on} k_{in}}{\beta(k_{on} + k_{off})k_{out}} \tag{S22}$$

We then derive the second-order partial derivatives, adding together the equations for the two gene states:

$$\frac{d(\langle n^2\rangle - \langle n\rangle)}{dt} = 2\alpha\langle n\rangle^{on} - 2(\beta + k_{in})(\langle n^2\rangle - \langle n\rangle) + 2k_{out}\langle nm\rangle \tag{S23}$$

$$\frac{d\langle nm\rangle}{dt} = \alpha\langle m\rangle^{on} + k_{in}(\langle n^2\rangle - \langle n\rangle) + k_{out}(\langle m^2\rangle - \langle m\rangle) - (\beta + k_{in} + k_{out})\langle nm\rangle \tag{S24}$$

$$\frac{d(\langle m^2\rangle - \langle m\rangle)}{dt} = 2k_{in}\langle nm\rangle - 2k_{out}(\langle m^2\rangle - \langle m\rangle) \tag{S25}$$

We derive $\langle n\rangle^{on}$ and $\langle m\rangle^{on}$ at steady state, from the system of four equations defined by the first derivatives of equations S10 and S11. Then we solve the system of equations S23-S25 at steady state, for $(\langle n^2\rangle - \langle n\rangle)$ and $(\langle m^2\rangle - \langle m\rangle)$. The Fano factors are simply:

$$F_n = \frac{\langle n^2\rangle - \langle n\rangle^2}{\langle n\rangle} = 1 + \frac{\langle n^2\rangle - \langle n\rangle}{\langle n\rangle} - \langle n\rangle \tag{S26}$$

$$F_m = \frac{\langle m^2\rangle - \langle m\rangle^2}{\langle m\rangle} = 1 + \frac{\langle m^2\rangle - \langle m\rangle}{\langle m\rangle} - \langle m\rangle \tag{S27}$$

Defining the variables $\gamma = k_{on} + k_{off}$ and $\delta = \beta + k_{in} + k_{out}$, the Fano factors are:

$$F_n = 1 + \frac{\alpha k_{off}[(\beta + k_{out})(\gamma + k_{out}) + k_{in}k_{out}]}{\gamma\delta[\gamma(\gamma+\delta) + \beta k_{out}]} \tag{S28}$$

$$F_m = 1 + \frac{\alpha k_{in}k_{off}(\gamma+\delta)}{\gamma\delta[\gamma(\gamma+\delta) + \beta k_{out}]}. \tag{S29}$$

#### Extended model with condensate-associated mRNA degradation

Adding the condensate-associated degradation reaction $(n, m)^{on/off} \to (n, m-1)^{on/off}$ to the master equations, with propensity $mk_{deg}$, and transforming them using the same generating functions (eq. S9), we derive:

$$\frac{\partial G^{on}}{\partial t} = k_{on}G^{off} - k_{off}G^{on} + \alpha(x-1)G^{on} + \beta(1-x)\frac{\partial G^{on}}{\partial x} + k_{out}(x-y)\frac{\partial G^{on}}{\partial y} + k_{in}(y-x)\frac{\partial G^{on}}{\partial x} + k_{deg}(1-y)\frac{\partial G^{on}}{\partial y} \tag{S30}$$

$$\frac{\partial G^{off}}{\partial t} = k_{off}G^{on} - k_{on}G^{off} + \beta(1-x)\frac{\partial G^{off}}{\partial x} + k_{out}(x-y)\frac{\partial G^{off}}{\partial y} + k_{in}(y-x)\frac{\partial G^{off}}{\partial x} + k_{deg}(1-y)\frac{\partial G^{off}}{\partial y} \tag{S31}$$

Following the same procedure as before, we derive analytical expressions for $\langle n\rangle$, $\langle m\rangle$, and $F_n$:

$$\langle n\rangle = \frac{\alpha k_{on}(k_{deg} + k_{out})}{(\beta k_{deg} + \beta k_{out} + k_{in}k_{deg})(k_{on} + k_{off})} \tag{S32}$$

$$\langle m\rangle = \frac{k_{in}}{(k_{deg} + k_{out})}\langle n\rangle = \frac{\alpha k_{on}k_{in}}{(\beta k_{deg} + \beta k_{out} + k_{in}k_{deg})(k_{on} + k_{off})} \tag{S33}$$

$$F_n = 1 + \frac{\alpha}{\mu\xi}\left[\frac{RS + k_{in}k_{out}T\xi}{\nu L} - \frac{k_{on}\xi^2}{\gamma}\right] \tag{S34}$$

where:

$$\gamma = k_{on} + k_{off} \qquad \xi = k_{deg} + k_{out} \qquad \mu = k_{deg}k_{in} + \beta\xi \qquad \nu = \beta + k_{in} + \xi \tag{S35}$$

$$L = (k_{deg} + \gamma)(\beta + k_{in} + \gamma) + (\beta + \gamma)k_{out} \tag{S36}$$

$$R = k_{deg}(\beta + k_{in} + k_{deg}) + k_{out}(\beta + 2k_{deg}) + k_{out}^2 \tag{S37}$$

$$S = k_{deg}(k_{in} + k_{on})(k_{deg} + \gamma) + k_{on}k_{out}(k_{in} + \gamma) + k_{deg}k_{out}(k_{in} + 2k_{on}) + k_{on}k_{out}^2 + \beta\xi(k_{deg} + k_{out} + \gamma) \tag{S38}$$

$$T = k_{deg}(k_{in} + k_{on}) + \beta(k_{deg} + k_{on} + k_{out}) + k_{on}(k_{in} + k_{out} + \gamma) \tag{S39}$$

### Monte Carlo Computer Simulations

#### Extended Model without Condensate-Associated mRNA Degradation

The first part of our stochastic simulations was performed to validate our analytical predictions for $\langle n \rangle$, $\langle m \rangle$, $F_n$, and $F_m$. In all cases, we varied $k_{in}$ along the x-axis as an effective measure of the level of condensation in the system. Specifically, we constructed curves $\langle n \rangle(k_{in})$, $\langle m \rangle(k_{in})$, $F_n(k_{in})$, and $F_m(k_{in})$ for three values of $\alpha$, investigating the behavior of the system for different mean burst sizes, and three values of $k_{off}$, investigating the behavior of the system for different mean burst durations. The complete sets of kinetic rates were chosen to produce mean nucleoplasmic mRNA copy numbers of 5, 20 and 50, respectively, as presented in Table S1.

**Table S1: No Degradation Parameter Values**

| Investigation with a set of $\alpha$ values | | Investigation with a set of $k_{off}$ values | |
|---|---|---|---|
| Simulation Steps | 10000000 | Simulation Steps | 10000000 |
| $\alpha$ | [12.5, 50, 125] $h^{-1}$ | $\alpha$ | 50 $h^{-1}$ |
| $\beta$ | 0.099 $h^{-1}$ | $\beta$ | 0.099 $h^{-1}$ |
| $k_{on}$ | 0.6 $h^{-1}$ | $k_{on}$ | 0.6 $h^{-1}$ |
| $k_{off}$ | 14.551 $h^{-1}$ | $k_{off}$ | [60, 14.551, 5.460] $h^{-1}$ |
| $k_{in}$ | (0-4) $h^{-1}$ | $k_{in}$ | (0-4) $h^{-1}$ |
| $k_{out}$ | 2 $h^{-1}$ | $k_{out}$ | 2 $h^{-1}$ |

The error bars in the simulations represent the standard errors of simulated values, calculated by block averaging. To derive the errors, we first performed an investigation by partitioning the trajectories into a number of equal time blocks and estimating the error

$$\sigma = \frac{\sigma_b}{\sqrt{N_b}} \tag{S40}$$

where $\sigma_b$ is the standard deviation of the block-means and $N_b$ is the number of blocks. This process aims to represent our trajectory as a number of independent measurements, which it achieves once $\sigma$ reaches a plateau at intermediate values of $N_b$. Then, we selected three values of $N_b = [30, 60, 100]$ to derive $\sigma$ for our simulations, selecting the largest $\sigma$ value as a conservative estimate of the standard error of the simulated values.

#### Extended Model with Condensate-Associated mRNA Degradation

We performed stochastic simulations to validate our analytical predictions for the extended model with condensate-associated mRNA degradation. Specifically, we constructed the curves $\langle n \rangle(k_{in})$, $\langle m \rangle(k_{in})$ and $F_n(k_{in})$, for three values of $k_{deg}$. The complete set of kinetic rates used is shown in Table S2, and the resulting curves in Figure S1.

**Table S2: Degradation Parameter Values**

| Investigation with a set of $k_{deg}$ values | |
|---|---|
| Simulation Steps | 10000000 |
| $\alpha$ | 50 $h^{-1}$ |
| $\beta$ | 0.099 $h^{-1}$ |
| $k_{on}$ | 0.6 $h^{-1}$ |
| $k_{off}$ | 14.551 $h^{-1}$ |
| $k_{in}$ | (0-4) $h^{-1}$ |
| $k_{out}$ | 2 $h^{-1}$ |
| $k_{deg}$ | [0.0, 0.1, 0.2] $h^{-1}$ |

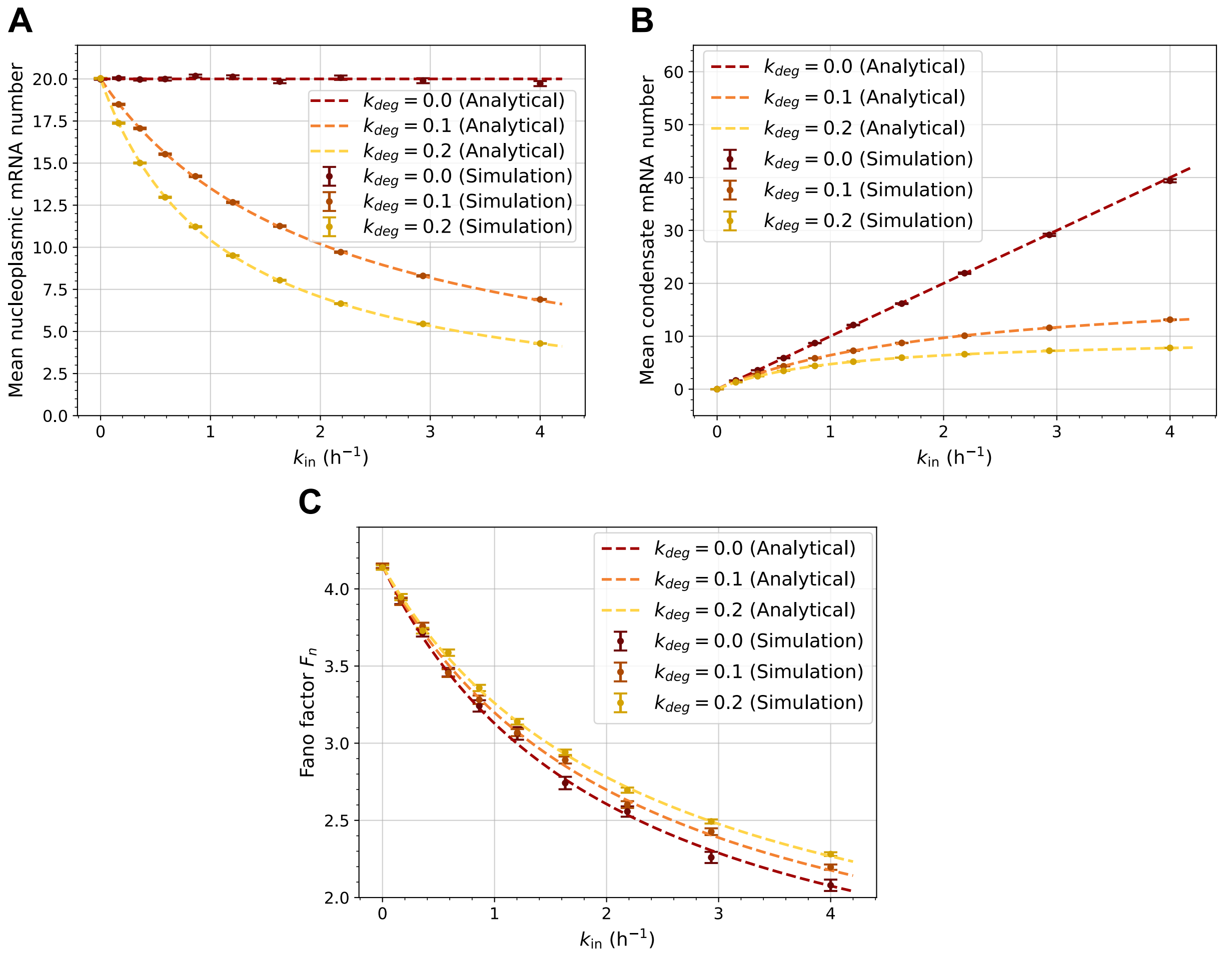


Figure S1: Mean numbers of mRNA molecules in different phases of the nucleoplasm and degree of nucleoplasmic mRNA stochastic fluctuations expressed via Fano factors for the extended model with condensate-associated degradation. A) Mean number of free mRNA molecules $\langle n \rangle$ for different condensate-associated mRNA degradation rates. B) Mean number of condensed mRNA molecules $\langle m \rangle$ for different condensate-associated mRNA degradation rates. C) Fano factor of free mRNA molecules, $F_n$, for different condensate-associated mRNA degradation rates. In all plots, $k_{in}$ is treated as an effective measure of mRNA condensation and the error bars represent the standard errors of the simulated values, calculated by block averaging.

Including condensate-associated mRNA degradation in the model, we observe a drastic change in the mean mRNA number in the condensates and the nucleoplasm, while the effects on stochastic noise (Fano factor) even for large $k_{deg}$ values are minimal. This justifies our approach to exclude condensate-associated mRNA degradation in our investigation of stochastic noise in the study and shows that regulation of transcriptional noise by condensates is achieved even with condensate-associated mRNA degradation.

#### Mean First-Passage Time and Effective Potential Simulations

To investigate the robustness of mRNA transcription in the presence of condensates, we performed simulations to estimate the mean first-passage time to return to the stationary state after a deviation in the nucleoplasmic mRNA number and the effective potential from the stationary probability distribution. For the mean first-passage time simulations, we performed simulations initiating the model in the state $[\langle n \rangle + x, \langle m \rangle]$, using $x = \pm 2, \pm 4$ for a range of $k_{in}$ values. The initial gene state was selected at random, based on the probability that the system is in the ON or OFF state (eq. S19). Thus, we assume that the deviation occurs at $t = 0$, independent of the gene state at that time. For the effective potential, we simulated trajectories for different values of $k_{in}$ to derive their stationary probability distribution. The complete sets of kinetic rates are shown in Table S3.

**Table S3: Mean First-Passage Time and Effective Potential Parameter Values**

| Mean First-Passage Time | | Effective Potential | |
|---|---|---|---|
| Number of Runs | 10000000 | Simulation Steps | 10000000 |
| $\alpha$ | 50 $h^{-1}$ | $\alpha$ | 50 $h^{-1}$ |
| $\beta$ | 0.099 $h^{-1}$ | $\beta$ | 0.099 $h^{-1}$ |
| $k_{on}$ | 0.6 $h^{-1}$ | $k_{on}$ | 0.6 $h^{-1}$ |
| $k_{off}$ | 14.551 $h^{-1}$ | $k_{off}$ | 14.551 $h^{-1}$ |
| $k_{in}$ | (0-4) $h^{-1}$ | $k_{in}$ | [0, 0.5, 2, 8, 40, 400] $h^{-1}$ |
| $k_{out}$ | 2 $h^{-1}$ | $k_{out}$ | 2 $h^{-1}$ |

The standard errors of simulated values were calculated once again by block averaging for the trajectories and propagation of error for the potential. Since the mean first-passage time trajectories were independent, the standard error of the mean value was used instead.

### Code

All code used in this study is publicly available in a GitHub repository:

https://github.com/alanit01/Condensate-Transcription
